# An evolutionary leap in plant reproduction: From archegonia and motile sperm to embryo sacs and pollen tubes

**DOI:** 10.1101/2025.08.04.668594

**Authors:** Dan Chen, Liwen Tang, Feng Wang, Mengxia Zhang, Siyuan Zhen, Meng Xia, Qian Wu, Mengxing Cao, Pengfei Jia, Yalong Guo, Fei Chen, Xiongbo Peng, Jin Hu, Mengxiang Sun, Hongju Li, Liangsheng Zhang, Weicai Yang

## Abstract

The transition of plants from water to land marks a pivotal step in their evolutionary history, with the change in reproductive mode being the most fundamental. This change freed plant reproduction from aquatic dependence, fostering terrestrial life’s prosperity. Key to the shift from archegonial to embryo sac reproduction are pollen tube emergence and female gametophyte changes, though their evolutionary trajectories and mechanisms remain unclear. *Ginkgo*, a basal gymnosperm, occupies a pivotal position in this transition, retaining motile sperm, simplified archegonia, and primitive pollen tubes, making it an ideal model. To study how the transition between these two reproductive modes occurred, we collected transcriptome data from the reproductive organs of Ginkgo and evolutionarily representative species. Our findings reveal the MYB98-CRP-ECS module- critical for angiosperm pollen tube guidance—in *Ginkgo*’s mature archegonia, with pollen tubes containing guidance components. While egg cells are transcriptionally and functionally conserved across land plants, changes in the cell fate of other female gametophyte component cells drove the shift from archegonia to embryo sacs. Pollen tubes may originate from fern male gametophyte internalization and modification. These findings clarify the evolutionary path and molecular basis of this major reproductive transition.

## Background

The evolution of plants from water to land represents a pivotal event in life history, accompanied by a series of morphological and physiological adaptations. Among these, the evolution of reproductive organs and associated shifts in reproductive strategies were critical drivers of this transition^1^. This evolutionary shift helped plants eliminate the limits of aquatic environments, enhanced their adaptability to dry land, and profoundly promoted the diversification of terrestrial life. However, the evolutionary steps and underlying mechanisms bridging the transition from archegonial to embryo sac reproduction remain poorly understood.

Land plants exhibit two primary reproductive modes: archegonial reproduction and embryo sac reproduction. Mosses and ferns rely on archegonial reproduction, where motile sperm (released into water by male structures) swim to reach female reproductive organs called archegonia for fertilization^2,3^. In contrast, flowering plants employ embryo sac reproduction, in which non-motile sperm are delivered to female structures (embryo sacs) via pollen tubes (Fig. 1A) ^4^. Over the past two decades, extensive research has explored pollen tube guidance mechanisms in angiosperms, making this topic a major focus in plant reproduction. Yet, such studies are largely restricted to angiosperms—predominantly the dicotyledonous Brassicaceae species *Arabidopsis thaliana*. In *Arabidopsis*, synergid cells (located at the ovule entry site for pollen tubes) express the MYB98(MYB domain protein 98)-CRP(cysteine-rich peptide)-ECS (Egg Cell-Secreted protein) module to guide pollen tubes, and various pollen tube components involved in this process have also been characterized^5,6^. Nevertheless, the origin and evolution of these guidance mechanisms, from sperm guidance to pollen tube guidance, is still remain unclear.

**Fig. 1.**
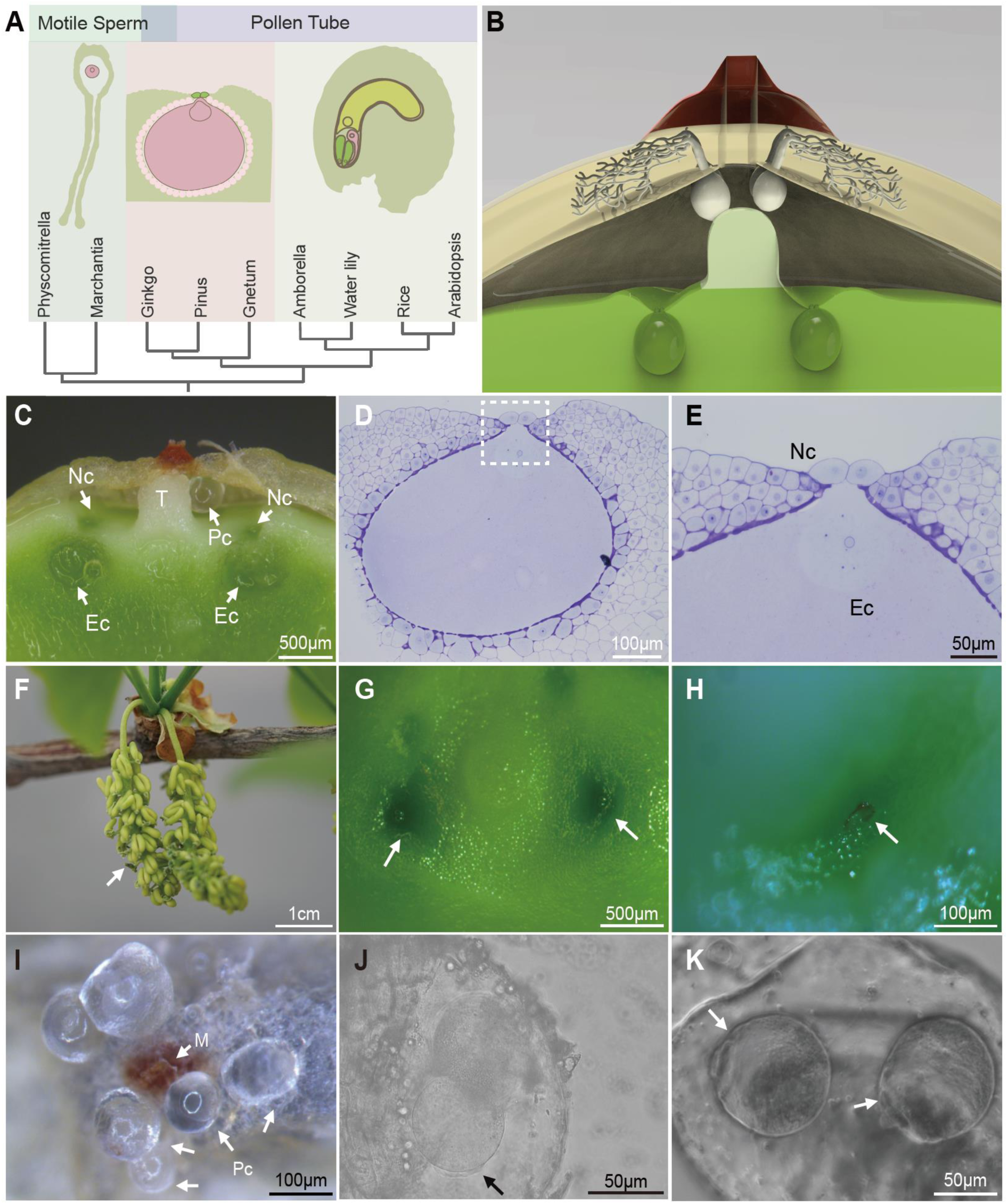
Evolution of female reproductive organs and fertilization dynamics in Ginkgo. **a**, Evolution of female reproductive organs. The pink cell represents the egg cell. **b,** Model of the reproductive structure of *Ginkgo biloba*. After micropylar absorption, branched pollen tubes grow slowly into ovular epidermal cells, with the pollen tube chamber suspended near the tentpole. Mature sperm form post neck cell genesis, swimming short distances through neck cells to reach the egg cell. **c**, Transection section of a mature archegonium. T, telepole. PC, pollen tube chamber. NC, neck cell. EC, egg cell. **d**, Semithin section of a mature archegonium. **e**, Enlarged view of the area outlined by the dashed box in **d**, highlighting neck cells (NC) and egg cell (EC). **f**, Mature microstrobilus (early April, Beijing). Arrowhead indicates an empty microsporangium post-pollen release. **g–h**, Ovules before **g** and after **h** fertilization. In **h**, a residual structure near the neck cell (arrowhead, August 23rd) suggested pollen tube rupture and sperm discharge. **i**, Saccate pollen tube chamber near micropyle. PC, pollen tube chamber. M, micropyle. **j**, Live-cell imaging of a dividing generative cell within a pollen tube chamber. The arrow indicates flagella. **k**, Two post-division sperm cells with motile flagella (arrows).

In gymnosperms, neck cells are reported to localize at the sperm entry site of archegonia. Their formation is temporally tightly linked to sperm development, and they degrade immediately after fertilization^7,8^. However, their role in sperm guidance and potential connections to angiosperm synergid cells have not been further investigated. The cellular evolutionary pathways between archegonia and embryo sacs, and the potential molecular genetic links between sperm guidance mechanisms and pollen tube guidance mechanisms, remain unclear.

To address these questions, we employed the basal gymnosperm *Ginkgo biloba*— a model organism that retains ancestral traits (motile sperm, archegonia) while possessing derived features (pollen tubes) ^9,10^—to investigate this reproductive transition. By integrating morphological and molecular data from *Ginkgo* with bioinformatic comparisons across evolutionarily key species, we aim to clarify the evolutionary pathway from archegonial to embryo sac reproduction, delineate the cellular evolutionary trajectory of critical reproductive cells, and elucidate the origins of core molecular modules governing pollen tube guidance. The findings of this study will significantly advance our understanding of the evolutionary principles underlying plant reproductive cells and refine plant evolutionary theory.

## Results

### The reproductive process of Ginkgo is characterized by two reproductive modes

Ginkgo exhibits two integrated reproductive modes, with dioecious characteristics. In Beijing, male microstrobilus mature in early April (Fig. 1F), while archegonia mature (Fig. 1C‒E) and fertilize (Fig. 1G‒H) in late August. Dry pollen grains are spindle-shaped (Fig. S1A), and hydrated ones are spherical with three visible nuclei and one indistinct degraded nucleus (Fig. S1B). Mature pollen tube chambers are suspended on the nucellus near the tentpole (Fig. 1H); each ovule contains 1–6 archegonia (most commonly two) and accommodates 1–9 pollen tube chambers.

In late August, 70–80% of ovules in individual trees fertilize within 2–3 days, with full fertilization completed in ∼5 days. Similar to Arabidopsis, the egg cell nucleus in mature archegonia migrates toward neck cells near the sperm entrance (Fig. 1D); sperm enter via the neck cell opening, which then degrades (Fig. S2). During peak fertilization, pollen tube chambers in an ovule vary in size, with only ∼1% containing two large motile sperm (Fig. 1I, K); most have undivided spermatogonia or ruptured chambers with liquid and a transparent shell near neck cells (Fig. 1H). This combination of flagellated motile sperm and pollen tube structures makes Ginkgo a model for plant reproductive evolution.

Live imaging captured generative cell division in the pollen tube chamber: one generative cell divided into two sperm, with flagella extending from opposite poles (Fig. 1J). Flagella reached motile length within minutes, initiating swimming, and the entire process was completed in ∼5 minutes. Flagellar activity ceased shortly after formation, suggesting rapid rupture post-sperm formation may explain the rarity of motile sperm-containing chambers.

Post-fertilization, ovules show morphological changes: integument turns from yellow-green to yellow, nucellus desiccates within a few days even in cases where some pollen tube chambers remained intact, the transparent shell near neck cells darkens and disappears within ∼10 days, and a deeper depression forms at the fertilization site (Fig. 1H)—all reliable indicators of successful fertilization.

### Construction of a gene expression atlas of the archegonium and the evolution of female reproductive organs

#### Transcriptomic profiles distinguish reproductive and vegetative tissues across species

To identify genes involved in Ginkgo reproductive cell specification, we sequenced transcriptomes of archegonia at different stages (AG, F), mature pollen (P), and pollen tubes (PT1, PT2) (Fig. S1, Table S1). Sample correlations (Fig. S3) align with PCA results (Fig. S4), with archegonium samples showing high correlation but fertilization-stage samples exhibiting weak correlation (likely due to asynchronous fertilization). Among the 27,836 genes in the Ginkgo genome, 27,746 were expressed in our data (log₂ TPM>1). The number of genes expressed in archegonia decreased from AG1 to AG4 but increased from F0 to F2 after fertilization (data S2). To identify tissue-specific expression, we added public transcriptomic data for Ginkgo roots, stems, and leaves for comparison. Uniform manifold approximation and projection (UMAP) analysis revealed nine distinct clusters, clearly separating reproductive and vegetative tissues (Fig. 2A).

**Fig. 2.**
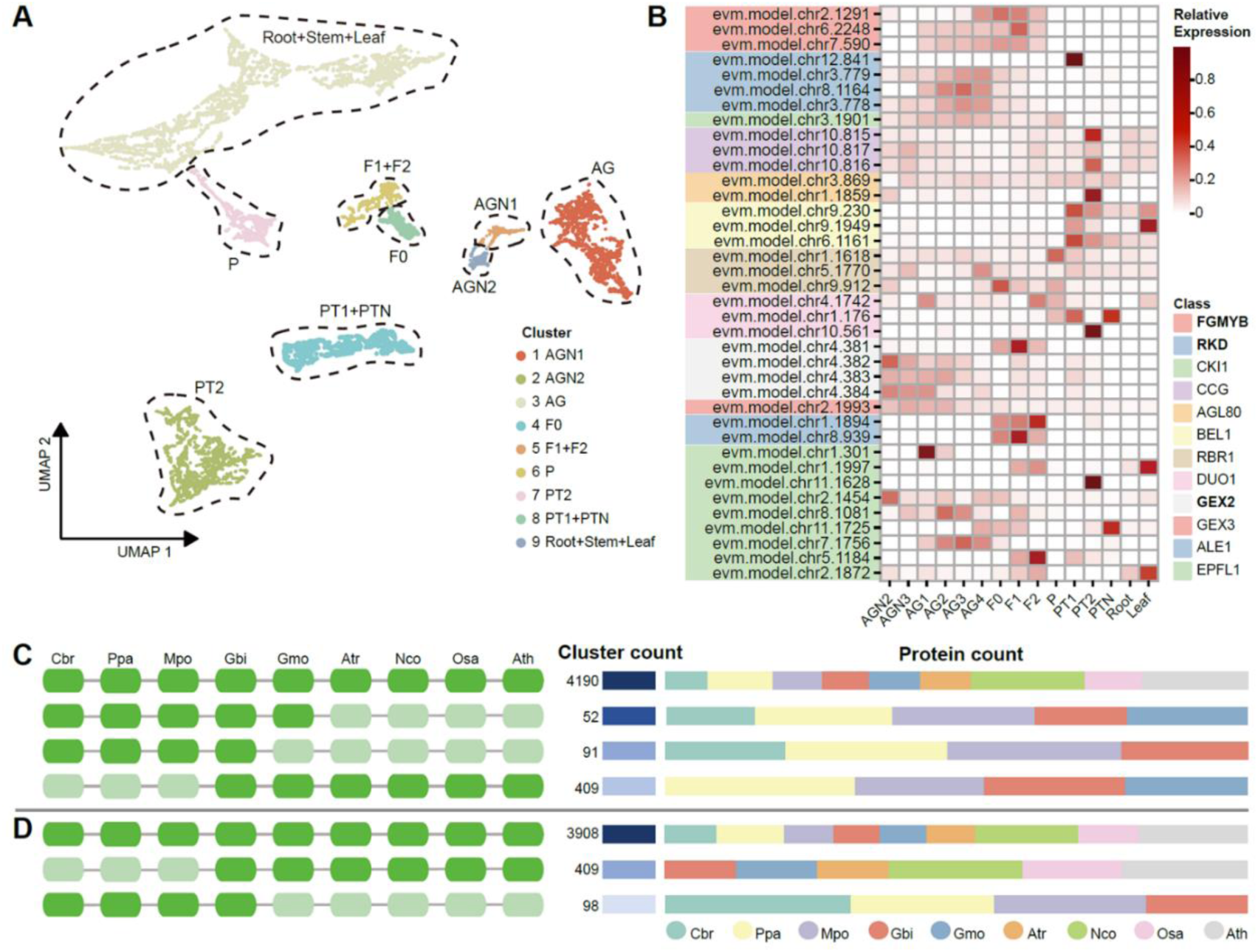
Transcriptomic features of Ginkgo reproductive development. **a**, UMAP plot of transcriptomic profiles from various Ginkgo tissues and developmental stages. Nine transcriptionally distinct clusters were identified, each representing a specific tissue type or developmental stage. **b**, Heatmap of gene expression levels for archegonium-specific expressed genes across different developmental stages. **c**, Shared orthogroups among nine representative plant species based on genes expressed in the Ginkgo archegonium (AG4), illustrating the conservation and divergence of gene families related to female gametophyte development. **d**, Shared orthogroups among nine representative plant species based on genes expressed in the growing Ginkgo pollen tube (PT2), illustrating the conservation and divergence of gene families related to pollen tube structure and function.

To explore conserved gene regulatory networks in gametophyte development, we analyzed representative plant species spanning key reproductive innovations: unicellular alga *Chara braunii*; archegoniate bryophytes such as *Marchantia polymorpha*^11^ and *Physcomitrella patens*^12^; gymnosperm *Gnetum montanum*^13^ (lacks archegonia); basal angiosperm *Amborella trichopoda*^14^ (eight-celled embryo sac); *Nymphaea colorata*^15^ (reduced four-nucleus embryo sac); and monocot/dicot flowering plants *Oryza sativa*^16^ and *Arabidopsis thaliana*^17^. *Ginkgo biloba*^18^ was used as the focal species. Selected public datasets (e.g., *Marchantia*^19^, *Ginkgo*^20^*, Cycas*^21^*, Amborella*^22^*, Nymphaea*^15^, *Oryza*) ^23^, together with newly generated transcriptomic datasets for key reproductive organs, were analyzed (data S6).

#### Conserved transcription factors and orthogroups reflect reproductive transitions

Systematic analysis of 1922 *Arabidopsis* transcription factors^24^ and reproductive regulators^5,6^ revealed evolutionarily conserved, tissue-specific expression in reproductive organs (Fig. S5). *FGMYB* (Female gamete MYB, evm.model.chr2.1291) and *RKD* (RWP-RK domain-containing protein, evm.model.chr3.778) showed the highest conservation in female gametophyte development, with archegonium-specific expression in *Ginkgo* (Fig. 2B). Some *CRPs* were highly expressed in mature *Ginkgo* archegonia (data S1), suggesting roles in female reproduction.

In male gametophytes, *DUO1* (DUO POLLEN 1) was the most conserved gene in our analysis, with one homolog specifically expressed in developing pollen tubes (containing generative cells) (Fig. 2B)—consistent with its conserved role in sperm cell differentiation across land plants^25^. *GEX2* (GAMETE EXPRESSED 2, evm.model.chr4.381), a sperm plasma membrane protein involved in *Arabidopsis* gamete fusion^5^, was upregulated in *Ginkgo* archegonia 1–3 days post-fertilization (undetectable in pollen), indicating that spermatogonia in Ginkgo may not initiate fertilization-related transcriptional activation until sperm cell genesis and release. This validates cytological markers (small black dots, water-filled depressions) for fertilization.

Orthologous gene comparisons were performed using genes expressed in *Ginkgo* AG4 stage (log₂ TPM>1) across eight phylogenetically representative species. Results showed that most orthogroups were shared, with the water lily (basal angiosperm) exhibiting the most prominent expansion (Fig. 2C). Notably, 52 orthogroups were uniquely shared by archegoniate plants (including Gnetum, which has reduced archegonia^26^), potentially involved in archegonium development. Additionally, 409 orthogroups unique to seed plants and expressed in *Ginkgo* archegonia reflect genetic changes underlying the structural transition from archegonium to seed plant reproductive architecture.

### The FGMYB-CRP-ECS module related to pollen tube guidance exists in Ginkgo archegonia

#### The phylogenetic and expression patterns of FGMYB genes are conserved

In Arabidopsis, synergids express MYB98 to regulator the transcription of cysteine-rich peptides (CRPs), such as LURE1s, which act as the pollen tube attractants. After fertilization, LURE 1 is either degraded by the egg cell-secreted aspartic proteases ECS1/2 or inactivated by nitrosylation modification to block further pollen tube entry^5,6^.

The R2R3-MYB subfamily member *MpFGMYB* is specifically expressed in female plants of *Marchantia polymorpha* and controls female sexual determination^19^. In Arabidopsis, the ancestral *FGMYB* gene has diversified into seven paralogs^19^, with *MYB98* regulating synergid cell specification^27–29^ and controls the expression of secreted CRP attractants, which play a major role in pollen tube guidance^30–33^, MYB64 and MYB119 are involved in central cell differentiation^34^, and MYB115/MYB118 function in seed development^35^.

Phylogenetic analysis of full-length protein sequences from 312 species revealed a well-supported *FGMYB* clade across gymnosperms, basal angiosperms, and monocots (Fig. 3A), with distinct diversification patterns in gymnosperms and angiosperms. The *MYB98* clade is Brassicaceae-specific (Fig. S6), indicating potential neofunctionalization.

**Fig. 3.**
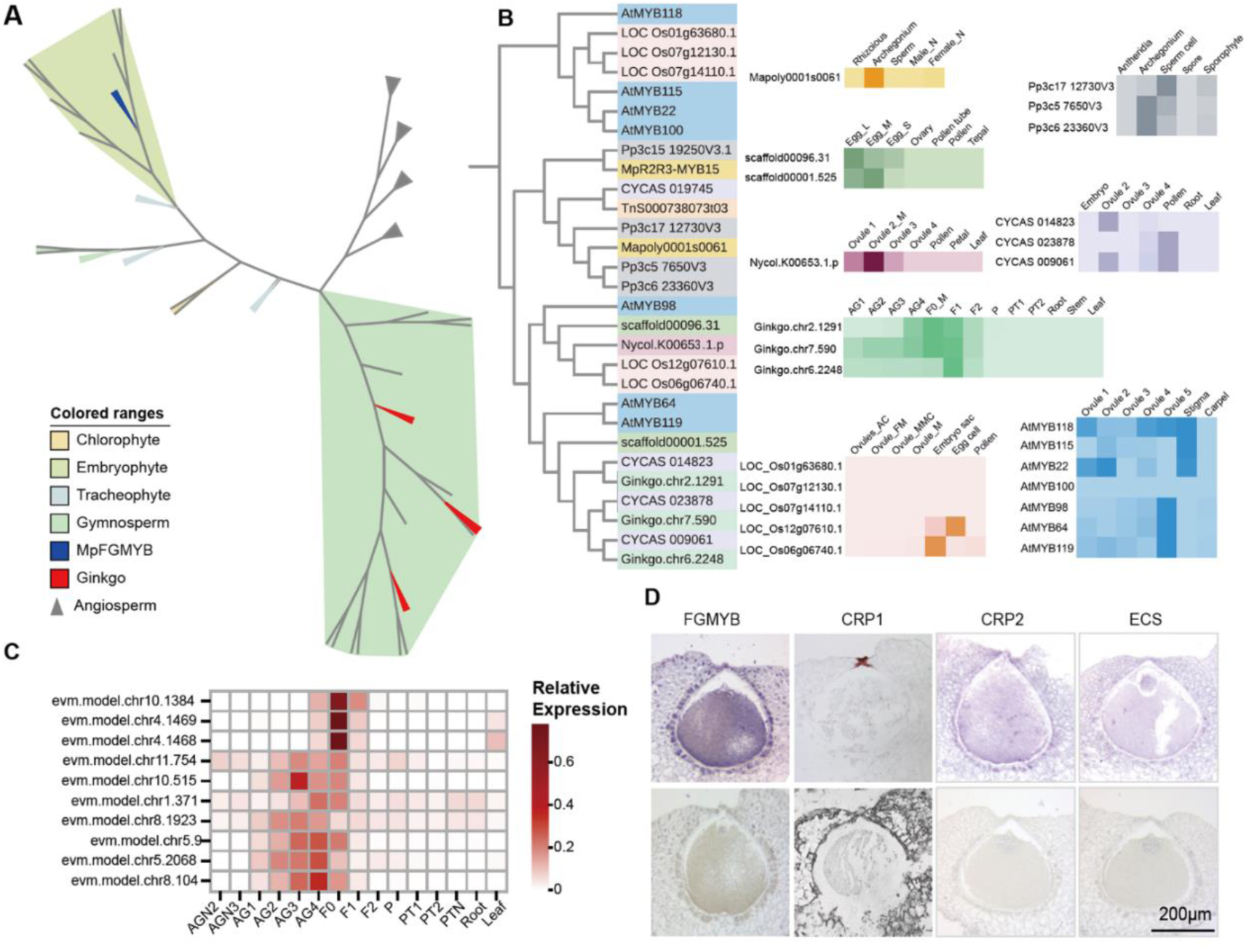
Expression patterns and phylogenetic analysis of FGMYB and related genes. **a**, Maximum-likelihood phylogenetic tree of FGMYB orthologs across 321 land plants (bootstrap support values greater than 70% are shown at major nodes). Species are color-coded as indicated in the legend. **b**, Phylogeny and expression patterns of FGMYB orthologs across eight representative plant species. The right panel presents a heatmap of transcript abundance (log₂ TPM) across reproductive and vegetative tissues, darker color indicates higher expression. **c**, Heatmap of the expression levels of ECSs and CRPs in Ginkgo, where darker color indicates higher expression. **d**, ISH of Ginkgo.chr2.1291 (FGMYB), evm.model.chr8.104 (CRP1), evm.model.chr4.1469 (CRP2), evm.model.chr10.1384 (ECS). The sense probe was used as a control and produced no obvious signals; scale bar = 200 μm. **Abbreviations**: Species: Mp, *Marchantia*; Pp, *P. patens*; sca, *A. trichopoda*; CYCAS, *C. panzhihuaensis*; Nycol, *N. colorata*; Ginkgo, *G. biloba*; LOC, *O. sativa*; At, *Arabidopsis*. Tissue types: Mp_Male_N: male vegetative organ; Mp_Female_N: female vegetative organ; Ginkgo_AG1–AG3: immature archegonia at different developmental stages; AG4: mature archegonium; F0: archegonium during fertilization; F1: archegonium after fertilization for 1–3 days; F2: archegonium for 7–10 days; pollen: mature pollen; petal: nonflower petals. Cycas_ovule 2: nonpollinated ovule; ovule_3/4: ovule 7/11 d post-pollination; Cycas_embryo: already fertilized embryo. sca_Egg_L: predominantly comprising egg cells; sca_Egg_M: enriched in young egg cells and third synergid cells; sca_Egg_S: predominantly consisting of synergid cells (transcriptome data from public sources). Nycol_ovule-od: the ovule of a bud one day before opening; Nycol_ovule-1d: the ovule of a bud just opening; Nycol_ovule-2d: the ovule of a bud 24 hours after opening; Nycol_ovule-3d: the ovule of a bud 48 hours after opening (petal and leaf transcriptome data from public sources). LOC_ovule_AC: AC stage ovule; ovules_FM: functional stage ovule; ovule_MMC: MMC stage ovule; ovule_M: mature ovule; Embryo sac: mature embryo sac (transcriptome data from public sources except embryo sac and pollen). At_ovule_1–5: different stages of the ovule (Ovule_5 collected during flowering stage; transcriptome data from public sources).

Expression analyses confirmed conserved roles in female reproductive tissues (Fig. 3B). In *Ginkgo*, three *FGMYB* homologs show stage-specific patterns: *Ginkgo.chr2.1291* peaks during fertilization, *Ginkgo.chr7.590* increases with archegonial development, and *Ginkgo.chr6.2248* rises post-fertilization (Fig. 3C). RNA in situ hybridization (ISH) localizes *Ginkgo.chr2.1291* to egg cells, jacket cells, and archegonium surfaces (Fig. 3D), resembling *Marchantia FGMYB* expression, supporting conserved roles in female gametophyte development across embryophytes.

Notably, the expression patterns of *FGMYB* homologs across species (Fig. 3B) further underscore their conserved association with female reproductive organ maturation and fertilization. For instance, in bryophytes, *MpFGMYB* in *Marchantia polymorpha* and two homologs in *Physcomitrella patens* are specifically expressed in archegonia^19^. Among seed plants, homologs in *Cycas* (gymnosperm), *Ginkgo biloba*, and *Nymphaea colorata* (basal angiosperm) reach peak expression during ovule maturation and decline post-fertilization. Similarly, *Arabidopsis thaliana MYB98* is highly expressed in mature ovules (predominantly in synergid cells) with reduced expression after fertilization, while homologs in *Oryza sativa* and *Amborella trichopoda* are specifically expressed in egg cells and mature embryo sacs. These consistent patterns across diverse plant lineages strongly support a conserved role of *FGMYB* genes in regulating female reproductive processes, particularly in coordinating archegonium/embryo sac maturation and fertilization.

#### CRP genes exhibit specific expression in archegonia

In *Arabidopsis*, pollen tube attractants are CRPs secreted by female gametophytic cells, and are transcriptionally regulated by *MYB98*^31,33^. In Ginkgo, 225 putative CRPs (E-value < 1e-5) were identified using HMMER models^36^ and InterProScan^37^ (data S1), with several archegonium-specific (Fig. 3C). *evm.model.chr8.104* (CRP1, DUF789 subfamily, Pfam: PF05623), showed the highest expression in mature archegonia, encoding a 280-amino acid peptide with four conserved cysteines and a 30-amino acid N-terminal signal peptide. ISH localized it to neck cells and adjacent jacket cells (Fig. 3D). We expressed the recombinant protein *in vitro* (Fig. S8B) and embedded it into agarose beads to treat *in vitro*–cultured pollen tubes (25-day cultures) and native pollen chambers; no pollen tube response was observed. Nevertheless, the specific localization and high expression of this gene in neck-associated cells suggest that it may function in neck cell differentiation or in facilitating sperm cell delivery during fertilization.

*evm.model.chr4.1469* (CRP2, CRP4820 subfamily) peaks at fertilization (F0), encoding a 70-amino acid peptide with nine conserved cysteines and a 24-amino acid N-terminal signal peptide. The ISH results revealed that this gene is expressed in egg cells, jacket cells, and cells adjacent to neck cells (Fig. 3D), implying a potential role in fertilization.

We classified all the CRPs into 441 subfamilies using the HMMER model^36^, only four of these subfamilies contained orthologs previously shown to be downstream targets of MYB98 in *Arabidopsis* (data S1). No conserved *MYB98*-target cis-elements were found in these archegonium-specific CRPs, indicating distinct regulatory mechanisms from *Arabidopsis*.

#### An ECS family member is upregulated post-fertilization in Ginkgo

Notably, an ECS family member, *evm.model.chr10.1384*, exhibited sharp and transient upregulation in archegonia after fertilization (Fig. 3C) with weak egg cell expression (Fig. 3D). Though not homologous to *Arabidopsis ECS1/2*^38^ (Fig. S7), its expression pattern resembles *ECS1/2* (which degrades LURE1 post-fertilization), suggesting conserved mechanisms across seed plants to remove attractant signals, restrict sperm guidance, and prevent polyspermy, which may be shared across seed plants to temporally restrict sperm guidance and prevent polyspermy.

### Evolutionary trajectory and model of female germ cells

#### RKD is phylogenetically conserved and specifically expressed in egg cells

*RKD* transcription factors act as evolutionarily conserved regulators of egg cell specification in land plants^39^. Comparative genomic and transcriptomic analyses across the nine representative species identified *RKD* as the most evolutionarily conserved gene among those related to egg cell specification and fertilization (Fig. 4A), with essential roles in both *Marchantia* and *Arabidopsis* (38). In *Ginkgo*, transcriptome analysis showed its *RKD* homolog is archegonium-specific (Fig. 4B), and ISH confirmed expression in egg cells and jacket cells (Fig. 4C).

**Fig. 4.**
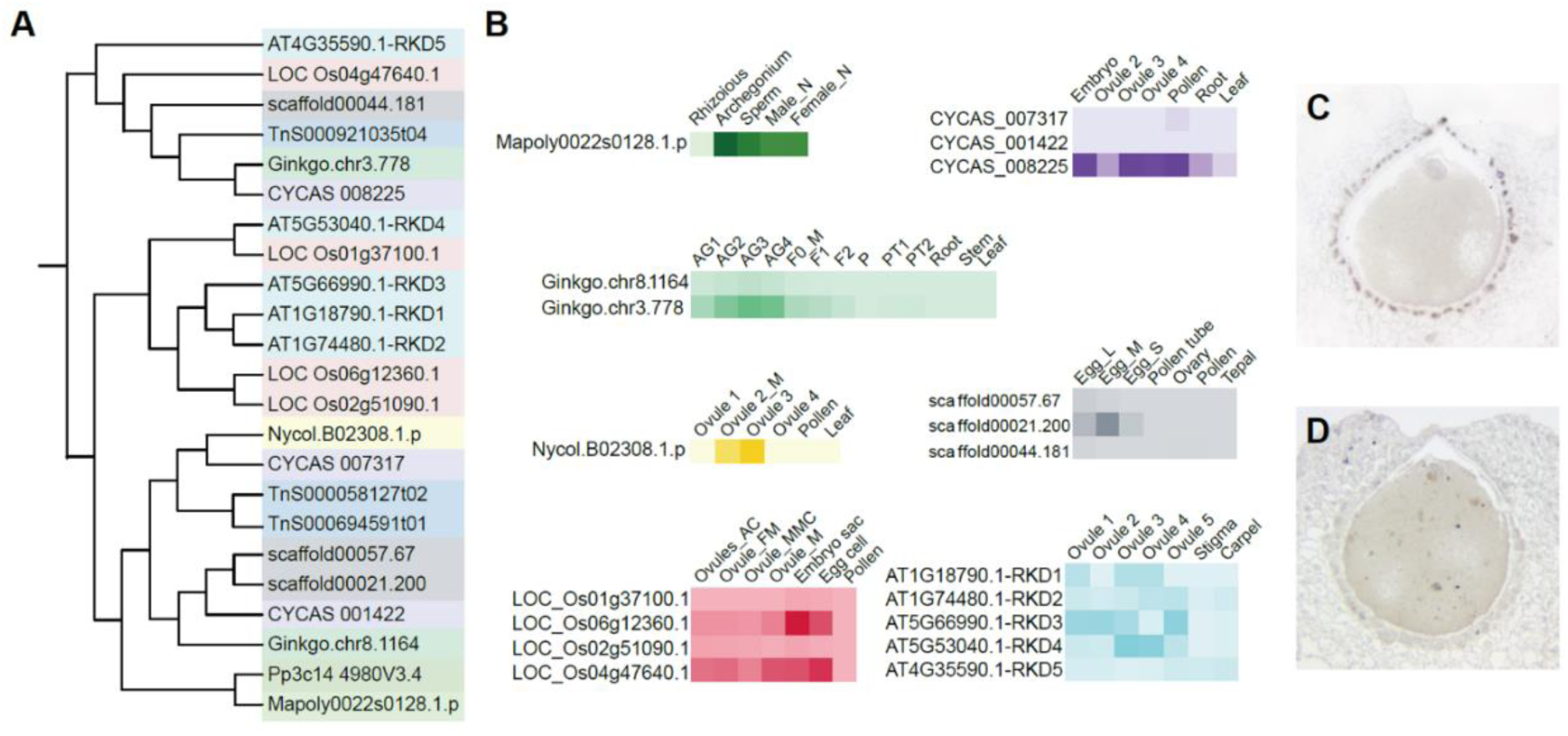
Conserved evolution and reproductive expression of RKD homologs in land plants. **a**, Phylogenetic tree of RKD homologs from nine representative land plant species, constructed using maximum likelihood methods based on full-length protein sequences. **b**, Heatmap of tissue-specific expression profiles of RKD homologs across seven representative species, with darker colors indicating higher expression. **c**, ISH of Ginkgo.chr3.778 in mature archegonia. **d**, Negative control using a sense probe, showing no detectable signal. The abbreviations are consistent with those defined in Fig. 5.

Cross-species expression patterns further support conservation: *Marchantia RKD* is most highly expressed in archegonia; *Cycas* homologs are expressed in late-stage ovules and pollen; *Amborella trichopoda* homolog *AmTr_v1.0_scaffold00021.200* is egg cell-specific (absent in synergids or vegetative tissues); rice homologs (*LOC_Os06g12360*, *LOC_Os04g47640*) are enriched in egg cells and mature embryo sacs; *Nymphaea colorata* homolog *Nycol.B02308.v1.2* peaks in mature ovules and 1 day post-fertilization, suggesting roles in egg specification and zygotic development (consistent with *Arabidopsis* functions). These patterns highlight the conserved expression of RKD in egg cell-adjacent cells across lineages.

#### Evolutionary model from the archegonium to the embryo sac

Based on gene-based phylogenetic trees, transcriptomic expression patterns, and ISH results, we propose an evolutionary model for the transition of female reproductive structures: Bryophyte and fern archegonia were simplified in gymnosperms (with shortened necks); the egg cell lineage remained transcriptionally and functionally conserved across land plants; gymnosperm neck/jacket cells likely evolved into angiosperm synergids; sister cells of the egg cell (persisting in late gymnosperm embryo sacs) gained fertilization competence, eventually forming angiosperm endosperm. This completes the transformation between reproductive modes (Fig. 1A), with *Ginkgo* representing a transitional state.

### Regulatory mechanisms by which *Arabidopsis* pollen tubes are activated in Ginkgo

#### Morphological observation and transcriptome acquisition of Ginkgo pollen tubes

The emergence of the pollen tube represents a key evolutionary innovation. As a basal gymnosperm, Ginkgo is the first known plant to possess a pollen tube structure, although it does not exhibit a guidance function. The branched pollen tube penetrates the inner integument of the ovule, with the pollen tube chamber remaining outside. Sperm cells form in the pollen tube chamber and are released shortly afterward. When the archegonium matures, the egg cells are ready, and the neck cells are fully developed. (Fig. 5A, B; Supplement video 1). Before fertilization, the inner integument contains an abundant fluid that degenerates after fertilization. Aniline blue staining showed callose accumulation in the distal region of 21-day in vitro-cultured pollen tubes (Fig. S8A), with in vivo tubes confined to the micropylar cavity of the inner integument (Fig. 5C, D).

**Fig. 5.**
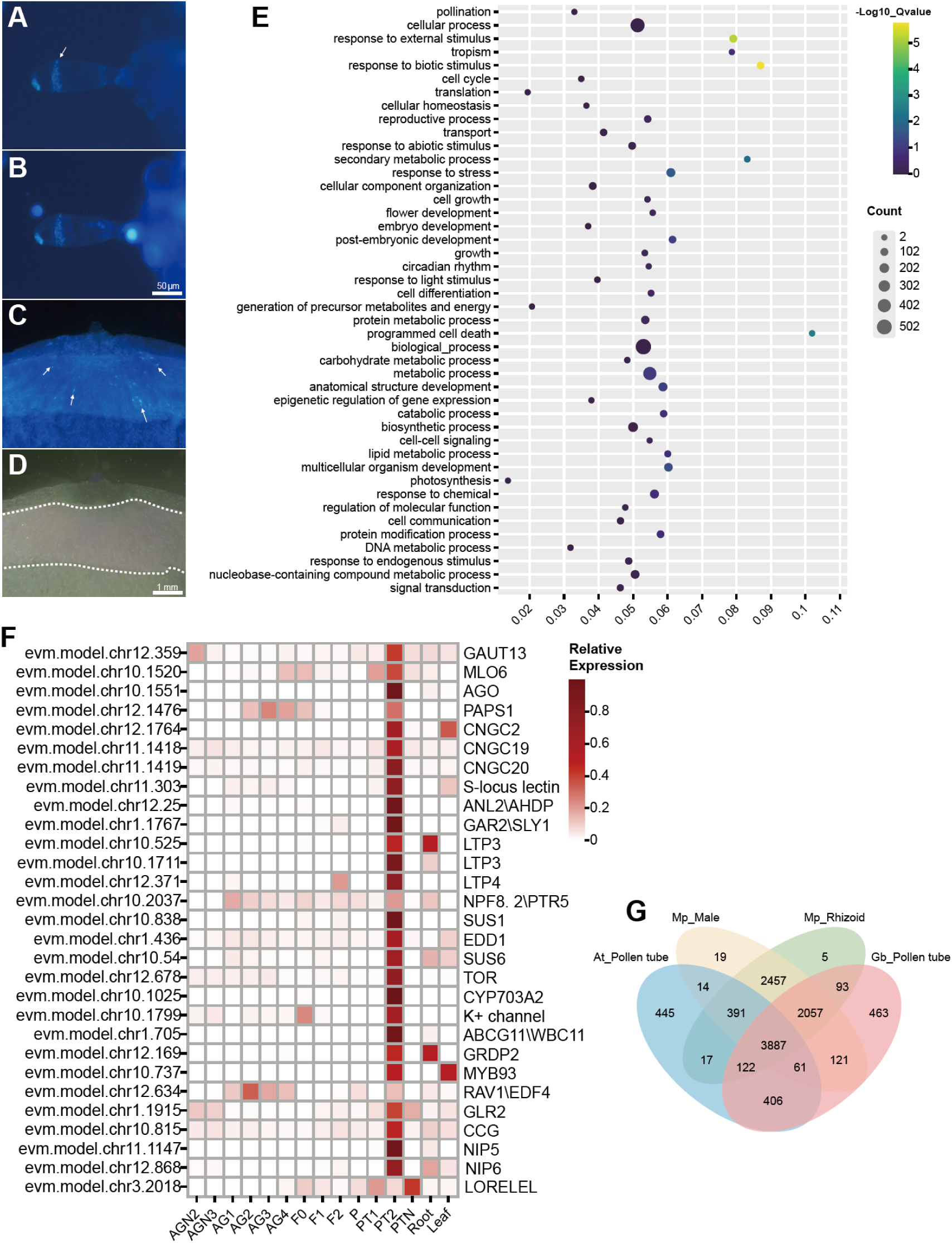
Evolution of the pollen tube structure in Ginkgo. **a**, Aniline blue staining of the pollen tube, highlighting the presence of callose in the distal region of the pollen tube. The arrow indicates the callose deposition. **b**, DAPI and aniline blue staining of the pollen tube. **c**, Aniline blue staining of the pollen tube within the endopleura of the ovule, illustrating pollen tube growth and callose deposition. **d**, Bright-field image of the pollen tube within the endopleura, with pollen tube growth restricted to the upper half (indicated by the dotted line). **e**, KEGG annotation of genes specifically expressed in the developing pollen tube (PT2), identified by UMAP analysis, highlighting genes involved in pollination and reproductive processes. **f**, Heatmap representation of gene expression related to pollen tube development, illustrating differential expression across various stages. **g**, Venn diagram of shared orthogroups for the pollen tube between *Ginkgo* and other plant species, demonstrating evolutionary divergence and similarity.

Transcriptome analysis included mature pollen, fertilization-stage pollen tubes (PT1, with inseparable inner integument tissue), and growing pollen tubes (PT2, collected 2 months pre-fertilization to avoid RNA degradation in aged PT1). PT1 expressed sperm formation genes but lacked guidance-related transcripts. This is likely because pollen tubes had stopped growing and developing before fertilization, so associated RNAs were no longer expressed, while PT2 provided abundant RNA for functional analysis.

#### Conserved molecular regulators involved in pollen tube growth and guidance in Ginkgo

To investigate the molecular basis of pollen tube formation in Ginkgo, we conducted Kyoto Encyclopedia of Genes and Genomes (KEGG) enrichment analysis on 1,913 PT2-specific genes (containing some unseparated inner integument of the ovule) identified from the UMAP analysis results. The enrichment analysis annotated six pollination-related genes and 78 reproduction-related genes (Fig. 5E, data S3), including key regulators of pollen tube formation, such as *GAUT13/14*, *LRE*, *LLG1*, *PAPS1*, and the pollen tube guidance genes *CNGCs*, *GLRs*, potassium channels, *MLO*, and *CCG*^5,6^ (Fig. 5H, data 4). In *Arabidopsis*, *GAUT13/14* expression in pollen tubes is essential; double mutation of these genes disrupts pectin distribution, causing growth defects. *LRE*, a GPI-anchored protein in synergid filiform apparatuses, prevents pollen tube overgrowth^40^. In Ginkgo, the specific expression of LRE in the apical integument (vs. lower integument) of PT2 suggests that LRE regulates tube growth restriction. *PAPS1*, which is involved in *Arabidopsis* tube penetration/elongation, has a PT2-specific homolog that may drive Ginkgo tube growth in the inner integument.

*CNGCs*, play crucial roles in pollen tube guidance in gymnosperm, three of which are expressed in PT2 (Fig. S9). Mirrors *the* roles in male gametophyte growth and *CNGC18* localization reported for Arabidopsis^41^, *MLO1* homolog in PT2 (lower in PT1). Six potassium channel genes suggest conserved ion signaling for tip growth (Fig. 5H, data 5). These ion channel-related genes suggest a conserved signaling network underlying pollen tube tip growth.

In addition to core pollen tube regulators, PT2 expresses homologs of genes critical for pollen development: *SPL8*, *PAPS1*, *GRDP2*, *PXL1*, *SGT*, *GAUT13*, *DAF*, and *CYP703A2*^42,43^. Flowering time regulators (*AGL6*, *CLPS3*, and *AP2*) ^43,44^ may be involved in regulating the synchronous maturation of archegonia and sperm in Ginkgo, reflecting complex reproductive gene networks (data 5).

#### Orthogroups analysis reveals that the Ginkgo pollen tube is most similar to liverwort male gametophytes

To investigate the origin of the pollen tube structure in *Ginkgo biloba*, we compared the transcriptomes of Ginkgo pollen tubes with those of *Marchantia* rhizoids, *Marchantia* male gametophytes, and *Arabidopsis* pollen tubes, with a focus on the evolutionary relationships of homologous genes. Venn diagram analysis revealed 3,887 shared orthogroups (conserved core genes); 2,057 orthogroups with *Marchantia* rhizoids and 2,457 with its male gametophytes; 463 *Ginkgo*-specific and 445 *Arabidopsis*-specific orthogroups (lineage innovations). Notably, *Ginkgo* pollen tubes shared the most orthogroups with *Marchantia* male gametophytes (Fig. 5G), suggesting a transitional evolutionary position—retaining conserved genetic links to early land plants while exhibiting unique innovations in pollen tube structure.

3,887 orthogroups were shared among all four tissues, suggesting a conserved core gene set. The Ginkgo pollen tube shared 2,057 orthogroups with *Marchantia* rhizoids and 2,457 with *Marchantia* male gametophytes, supporting partial transcriptomic similarity. Notably, 463 orthogroups were unique to the Ginkgo pollen tube, and 445 were specific to the *Arabidopsis* pollen tube, indicating lineage-specific innovations. Notably, *Ginkgo* pollen tubes shared the highest number of orthogroups with *Marchantia* male gametophytes (Fig. 5G), supporting the hypothesis that the pollen tube structure in *Ginkgo* likely originated from the rhizoids produced by male gametophytes in archegoniate plants. This evolutionary connection highlights a conserved genetic foundation linking early land plant reproductive structures to the specialized pollen tube in gymnosperms.

## Discussion

How land plants transitioned from archegonium-based reproduction to embryo sac–based reproduction remains a central question in plant evolutionary biology. Although the mechanisms of pollen tube guidance in angiosperms have been extensively studied, the evolutionary trajectory connecting these two reproductive modes remains poorly understood. Our morphological microscopic observations and molecular analyses of Ginkgo reproductive cells clarify the evolutionary path from archegonial to embryo sac reproduction, and identify the evolutionary trajectories of major gamete cells (neck cells evolving into synergids, with egg cells remaining conserved). Notably, our findings reveal that the core molecular module for pollen tube guidance has already been established in gymnosperms.

Although pollen tubes do not yet have a guidance function in Ginkgo, we identified orthologs of angiosperm pollen tube guidance genes, including the *MYB98-CRP-ECS* module. These genes exhibit highly consistent expression patterns during fertilization in Ginkgo and Arabidopsis, suggesting that the pollen tube guidance mechanism in flowering plants may have evolved from the sperm guidance mechanism. This conservation implies that the molecular framework for reproductive cell communication was co-opted and refined during the transition to flowering plants, rather than being a de novo innovation.

The conserved expression of *FGMYB* and *RKD* in egg cells and their adjacent cells across eight species further supports the deep conservation of female gametophyte regulatory modules in land plants. In *Ginkgo*, the most highly expressed archegonium-specific CRP (*evm.model.chr4.1469*) localizes specifically to neck cells, mirroring the synergid-specific expression of CRPs in *Arabidopsis*. This continuity in CRP expression patterns reinforces the hypothesis that synergids are derived from archegonial neck cells, with molecular functions retained despite morphological divergence.

Based on cellular expression patterns of these key genes, we propose a refined model for the evolution of female gametophytes during the archegonium-to-embryo sac transition. Egg cells, conserved across land plants, maintain their core function in fertilization and embryogenesis. Synergids, by contrast, likely arose from archegonial neck cells: in mosses and ferns, archegonia possess distinct flask-shaped structures with prominent necks; in gymnosperms, archegonia are internalized and enclosed, with neck regions morphologically reduced; in late-diverging gymnosperms such as *Gnetum*, egg cell sister cells persist in the mature gametophyte, gain fertilization competence^26^, and eventually give rise to the endosperm after double fertilization in angiosperms. This sequence clarifies the cellular evolutionary path underlying the transformation of female reproductive structures.

Our findings also shed light on the evolutionary process of the transition from motile-dependent to pollen tube-mediated fertilization. In mosses and ferns, free-living male gametophytes produce motile sperm for fertilization. In the gymnosperms Ginkgo and Cycas, male gametophytes develop within ovules: branched pollen tubes grow into the inner integument, leaving outside a pollen tube chamber containing spermatogonia. Rather than transporting sperm, the pollen tube chamber ruptures near the neck cells, releasing sperm that enter through the neck cell opening and fuse with the waiting egg nucleus, thereby completing fertilization. In angiosperms and most gymnosperms, nonmotile sperm are delivered directly to female gametophytes via pollen tubes.

This transition was accompanied by coordinated modifications in female gametophyte structure, reflecting a stepwise reuse of cellular and molecular modules— consistent with evolutionary strategies observed in animals. This suggests a fundamental principle of biological evolution: the conservation of core regulatory networks, with lineage-specific modifications enabling functional innovation.

## Methods

### Materials

Ginkgo biloba samples used in this study were collected from trees cultivated on the Olympic Village campus of the Chinese Academy of Sciences, Beijing. Detailed sampling dates for each developmental stage are provided in Table S1. On this campus, ovule fertilization typically initiates in early August, although inter-cultivar variation may result in a shift of up to one month. Even among closely related cultivars, fertilization timing can differ by as much as two weeks. For an individual tree, approximately 70% of ovules undergo fertilization within a two-day window, and full fertilization of all ovules is typically completed within 4 to 5 days. Each ovule contains one to six archegonia, with two being the most frequently observed.

Fresh female cones were quickly dissected under a microscope, four cuts were made around the archegonium and one horizontal cut, resulting in samples approximately 1.5 mm × 1.5 mm × 2 mm in size. Each sample consisted of 6-8 archegonia. Pollen grains were collected during the peak of pollen release in early spring when the male microstrobilus were slightly yellow, and part of the anthers had already released pollen. The pollen was allowed to naturally disperse on a petri dish at room temperature. Only high-quality pollen, which naturally dispersed, was used for RNA extraction and library construction. The pollen that was shaken out forcefully was of lower quality and was avoided.

At the peak fertilization period in August, mature pollen tube (PT1) chambers were collected. As pollen tube chambers are delicate and can rupture easily when removed, the tissue surrounding the pollen tube chamber was carefully dissected, with the inner integument (approximately 3 mm × 4 mm) containing the pollen tube chambers collected directly. Tissue surrounding the pollen tube chambers, devoid of pollen tubes, was used as a control.

By isolating archegonia at different developmental stages, we observed that the neck cells at the micropylar end exhibit no obvious depression in immature archegonia. As the archegonium matures, a distinct depression gradually forms at the neck region, and the color of the area surrounding the neck cells changes from a translucent pale yellow-green to an opaque green. Following fertilization, the integument begins to yellow within a few days.

### RNA Extraction and Quality Control

Total RNA was extracted using TRIzol™ Reagent (Invitrogen, 15596026) for most tissue samples. For RNA isolated from laser capture microdissection (LCM) materials, the PicoPure™ RNA Isolation Kit (Ambion, KIT0214) was employed. For Ginkgo pollen and leaf tissues, TRIzol™-based extraction was found to be suboptimal, likely due to high levels of polysaccharides and polyphenolic compounds, the RNA was extracted using the MiniBEST Plant RNA Extraction Kit (Takara, 9769) instead.

RNA integrity was initially checked using 0.8% agarose gel electrophoresis, and RNA concentration was determined using a NanoDrop 2000C Spectrophotometer. Samples were selected based on the presence of clear 25S rRNA and 18S rRNA bands, absence of the 5S rRNA band, and a concentration 100-200 ng/μL. The quality of the RNA was further assessed using the Agilent 2100, and samples with high RNA Integrity Number (RIN), low protein content, and minimal genomic DNA contamination were selected. These samples were then treated with DNase to remove any genomic DNA before being used for RNA-sequencing.

### RNA Sequencing and Library Construction

Due to the low RNA yield from Ginkgo seeds, even after increasing the sample size, conventional library construction remained insufficient. Thus, micro-quantity RNA sequencing was employed. Quality-controlled RNA samples were amplified using the Smart-seq2 protocol, followed by standard eukaryotic transcriptome library preparation. Sequencing was performed on the BGISEQ platform with a PE150 strategy, generating 6G clean data per sample (Q20 ≥ 80%). Individual samples met ≥90% of the target data volume when total data requirements were satisfied. Raw reads were processed using SOAPnuke to filter low-quality sequences, adapter contamination, and high-N-content reads. Clean reads were aligned to the reference genome using HISAT and Bowtie2 for new transcript prediction, SNP/InDel analysis, and differential splicing gene detection.

### Transcriptomic Data Analysis

Transcriptomic analysis utilized the 2019 updated *Ginkgo biloba* genome from GigaDB. Sequencing on the BGISEQ platform (PE150) yielded 6G clean data per sample, with individual samples meeting ≥90% of the target volume. DNBSEQ platform sequencing provided an average of 6.24 Gb/sample, with 91.66% genome alignment and 81.79% annotated gene set alignment. A total of 39,775 new genes were predicted, with 74,593 expressed genes (35,245 known; 39,348 novel). Additionally, 61,885 new transcripts were identified, including 6,311 novel splice variants, 39,775 new protein-coding transcripts, and 15,799 long non-coding RNAs. Data were analyzed for differentially expressed genes (DEGs), KEGG pathway enrichment, and visualized via UMAP and principal component analysis (PCA).

### In Situ Hybridization (ISH)

For ISH, RNA probes for Ginkgo.chr2.1291 (FGMYB), evm.model.chr8.104 (CRP1), evm.model.chr4.1469 (CRP2), and evm.model.chr10.1384 (ECS) were synthesized. The ISH was performed on paraffin-embedded sections of Ginkgo archegonia at various stages of development. A sense probe was used as a negative control, showing no detectable signal. The tissue sections were analyzed under a microscope, and images were captured to evaluate the spatial expression of these genes.

### Statistical Analysis

For RNA-seq data, gene expression levels were quantified using FPKM (Fragments Per Kilobase Million) values. Differential expression analysis was performed using the DESeq2 package, and genes with a fold change greater than 2 and a p-value less than 0.05 were considered significant. GO enrichment and KEGG pathway analysis were performed to identify functional categories and biological pathways associated with the differentially expressed genes.

### UMAP Analysis

After applying log₂ transformation and Z-score normalization to the expression data (FPKM values) of all genes, we assessed gene expression specificity using the tissue specificity index (TAU). This metric quantifies the degree of expression specificity by comparing gene expression distributions across multiple tissue samples. Genes with a TAU score greater than 0.6 were considered to exhibit high tissue specificity. These genes were then subjected to nonlinear dimensionality reduction using the UMAP algorithm (version 0.2.10.0), with the following parameters: n_neighbors = 7, n_epochs = 2000, and min_dist = 0.01. The resulting low-dimensional representations were visualized as two-dimensional scatter plots. Subsequently, genes were clustered using the DBSCAN algorithm to identify differentially expressed genes across distinct tissues.

### Orthologous gene analysis

Use OrthoFinder algorithm (E-value < 1e-5, inflation value = 1.5) on the OrthoVenn3 platform (https://orthovenn3.bioinfotoolkits.net).

### Note on Data Visualization

All figures, including line plots, heatmaps, and phylogenetic trees, were generated using the Chiplot software.

## Data availability

All the raw sequencing data for Ginkgo have been deposited at the NCBI and are accessible via the BioProject accession code PRJNA1274214.

## Acknowledgments

We thank Huijin Zhang (Institute of Botany, Chinese Academy of Sciences) for Nymphaea materials; Xiaorong Huang, Shan Liang, and Hui Zhou for ISH technical assistance; Yinjiao Xu for CRP analysis guidance; Jianguo Meng and Fei Yang for protein purification help; Yao Yao for semi-sectioning. This work was supported by: National Natural Science Foundation of China (32400295, 31991203); Research Startup Funding from Hainan Institute of Zhejiang University (0205-6602-A12201).

## Author contributions

D.C. conceptualized the study, developed methodology, performed investigation, created visualizations, administered the project, acquired funding, and wrote the original draft. W.C.Y. contributed to conceptualization, methodology, supervision, funding acquisition, and review and editing of the manuscript. S.Y.Z. and M.X. contributed to visualization. L.S.Z. contributed to methodology, and review and editing. H.J.L. contributed to methodology and review and editing. F.C. contributed to methodology, visualization, and review and editing. Y.L.G. contributed to methodology and review and editing. P.F.J., Q.W., and M.X.C. contributed to methodology. L.W.T. and F.W. contributed to investigation and visualization. M.X.Z., M.X.S., J.H., and X.B.P. contributed to review and editing.

## Materials & Correspondence

Correspondence and requests for materials should be addressed to.

